# Cytoplasmic ribosomes hitchhike on mitochondria to dendrites

**DOI:** 10.1101/2024.09.13.612863

**Authors:** Corbin J. Renken, Susie Kim, Youjun Wu, Marc Hammarlund, Shaul Yogev

## Abstract

Neurons rely on local protein synthesis to rapidly modify the proteome of neurites distant from the cell body. A prerequisite for local protein synthesis is the presence of ribosomes in the neurite, but the mechanisms of ribosome transport in neurons remain poorly defined. Here, we find that ribosomes hitchhike on mitochondria for their delivery to the dendrite of a sensory neuron in *C. elegans*. Ribosomes co-transport with dendritic mitochondria, and their association requires the atypical Rho GTPase MIRO-1. Disrupting mitochondrial transport prevents ribosomes from reaching the dendrite, whereas ectopic re-localization of mitochondria results in a concomitant re-localization of ribosomes, demonstrating that mitochondria are required and sufficient for instructing ribosome distribution in dendrites. Endolysosomal organelles that are involved in mRNA transport and translation can associate with mitochondria and ribosomes but do not play a significant role in ribosome transport. These results reveal a mechanism for dendritic ribosome delivery, which is a critical upstream requirement for local protein synthesis.

## INTRODUCTION

Neurons are highly polarized cells with functionally distinct compartments. The functional separation between axons and dendrites requires them to maintain distinct proteomes, often at great distances from the cell body. One mechanism employed by neurons to address this need is the synthesis of proteins directly “on site” in neurites through a process known as “local translation” (Holt et al., 2019). Local translation has been reported to play important roles in diverse cellular functions such as synaptic long-term potentiation (Kang and Schuman, 1996; Martin et al., 1997; Miller et al., 2002), growth cone guidance (Campbell and Holt, 2001; Leung et al., 2006), and axon regeneration (Yan et al., 2009; Terenzio et al., 2018); therefore, the upstream mechanisms that enable and regulate local translation are of great interest.

For local translation to occur, ribosomes must be present in the neurite. In the case of dendrites, the presence of ribosomes has been well-documented for many years (Bodian, 1965; Steward and Levy, 1982). The relative sparsity of ribosomes in dendrites compared to the cell body likely imposes a limiting factor on the overall capacity for dendritic protein synthesis (Ostroff et al., 2002; Sutton and Schuman, 2006), highlighting the importance of transport mechanisms that deliver ribosomes to the dendrite. However, the mechanisms regulating ribosome transport and positioning in dendrites remain poorly understood.

In the filamentous fungus *U. maydis*, ribosome transport is mediated by early endosomes (Higuchi et al., 2014). In neurons, most studies have focused primarily on the transport of mRNAs; however, early studies demonstrated that ribosomes can be found in messenger ribonucleoprotein (mRNP) granules (Knowles et al., 1996; Elvira et al., 2006), which can associate with kinesin and dynein motors or their adaptors (Kanai et al., 2004; Fukuda et al., 2020; Elvira et al., 2006; Baumann et al., 2022). Consistent with a role for microtubule motors in neuronal ribosome transport, a genetic screen in *C. elegans* found that mutations in tubulin cause ribosome misdistribution (Noma et al., 2017).

More recently, mRNAs in neurons have been observed co-transporting with membrane-bound organelles (Cioni et al., 2019; Liao et al., 2019; Harbauer et al., 2022; Cohen et al., 2022), raising the possibility that neuronal ribosomes might also be transported by organelle “hitchhiking” as in filamentous fungi. Whether such a mechanism plays a significant role in the transport of ribosomes to dendrites remains untested. An mRNA adaptor complex on Rab5-positive early endosomes partially colocalizes with mRNA in dendrites (Schuhmacher et al., 2023), and Rab7-positive late endosomes exhibit partial colocalization with translation “hotspots” containing mRNAs and ribosomes in axons (Cioni et al., 2019). However, dominant negative forms of Rab5 or Rab7 did not interfere with mRNA transport or localization (Cioni et al., 2019), suggesting that early and late endosomes are insufficient to instruct mRNA transport.

Lysosomes and mitochondria have also been implicated as mediators of RNA hitchhiking: precursor microRNAs (Corradi et al., 2020), microRNAs, and the RNAi machinery (Gershoni-Emek et al., 2018) all co-transport with lysosomes in the axon, as do mRNP granules (Liao et al., 2019), raising the possibility that lysosomes could transport ribosomes. Furthermore, two specific mRNAs were recently shown to transport on mitochondria in a translation-dependent manner (Harbauer et al., 2022; Cohen et al., 2022), suggesting that mitochondria might also serve as a vehicle for ribosome transport. This idea would be consistent with the close association between mitochondria and translationally active endosomes in axons (Cioni et al., 2019), as well as with the finding that lysosome-transported mRNAs can support mitochondrial health in the axon (De Pace et al., 2024). Nevertheless, ribosome transport *per se* – into axons or dendrites – has not been directly addressed.

Here we directly examine ribosome transport into the dendrites of sensory neurons in *C. elegans*. Leveraging the amenability of this model to live cell imaging, electron microscopy analysis and *in vivo* perturbations of organelle transport, we determined that 74% of motile dendritic ribosomes hitchhike on mitochondria. In an unbiased screen for ribosome localization we identified genes for mitochondria transport and morphology, and we employed genetic knockouts and gain-of-function manipulations to demonstrate that mitochondria are required and instructive for dendritic ribosome transport. We observed that ribosomes co-transport with distinct populations of mitochondria and that their association requires the atypical Rho GTPase MIRO-1. Examining several endolysosomal organelles, we find that RAB-7 is closely associated with mitochondria and ribosomes but does not instruct ribosome localization. These results demonstrate a role for mitochondria in dendritic ribosome transport and address a key mechanistic gap in our understanding of local translation.

## RESULTS

### Ribosomes are stereotypically positioned in the dendrites of AWB and AWC

To investigate the mechanisms of ribosome transport *in vivo*, we visualized the localization of ribosomes in the *C. elegans* chemosensory neurons AWB and AWC. Both neurons possess long dendrites that extend through the amphid sensilla, beginning from the soma in the nerve ring and terminating in large sensory cilia at the tip of the nose (Perkins et al., 1986) (Fig. 1 A). We chose to use these neurons both for their highly-visible dendrites and because previous work in the AWC dendrite has implicated a role for dendritic protein synthesis during olfactory adaptation (Kaye et al., 2009). To visualize ribosomes, we fused eGFP (F64L, S65T) (hereafter referred to as “GFP”) to the C-terminus of the 40S small ribosomal subunit protein RPS-2, a fusion construct that has previously been shown to assemble into ribosomes and complement a non-viable *rps2*Δ null allele in yeast (Milkereit et al., 2003) (Fig. 1 B). When expressed in AWB and AWC using the *odr-1p* promoter, RPS-2::GFP signal exhibits a pattern consistent with ribosome localization: it fills the cell body, enriching around the nucleus in a pattern reminiscent of the endoplasmic reticulum (Fig. 1 B). RPS-2::GFP can also be detected as discrete puncta within the nucleus, likely corresponding to nucleoli (Fig. 1 B). FRAP analysis of somatic signal revealed slow recovery of RPS-2::GFP after photobleaching, consistent with its incorporation into ribosomes (not shown).

**Figure 1.**
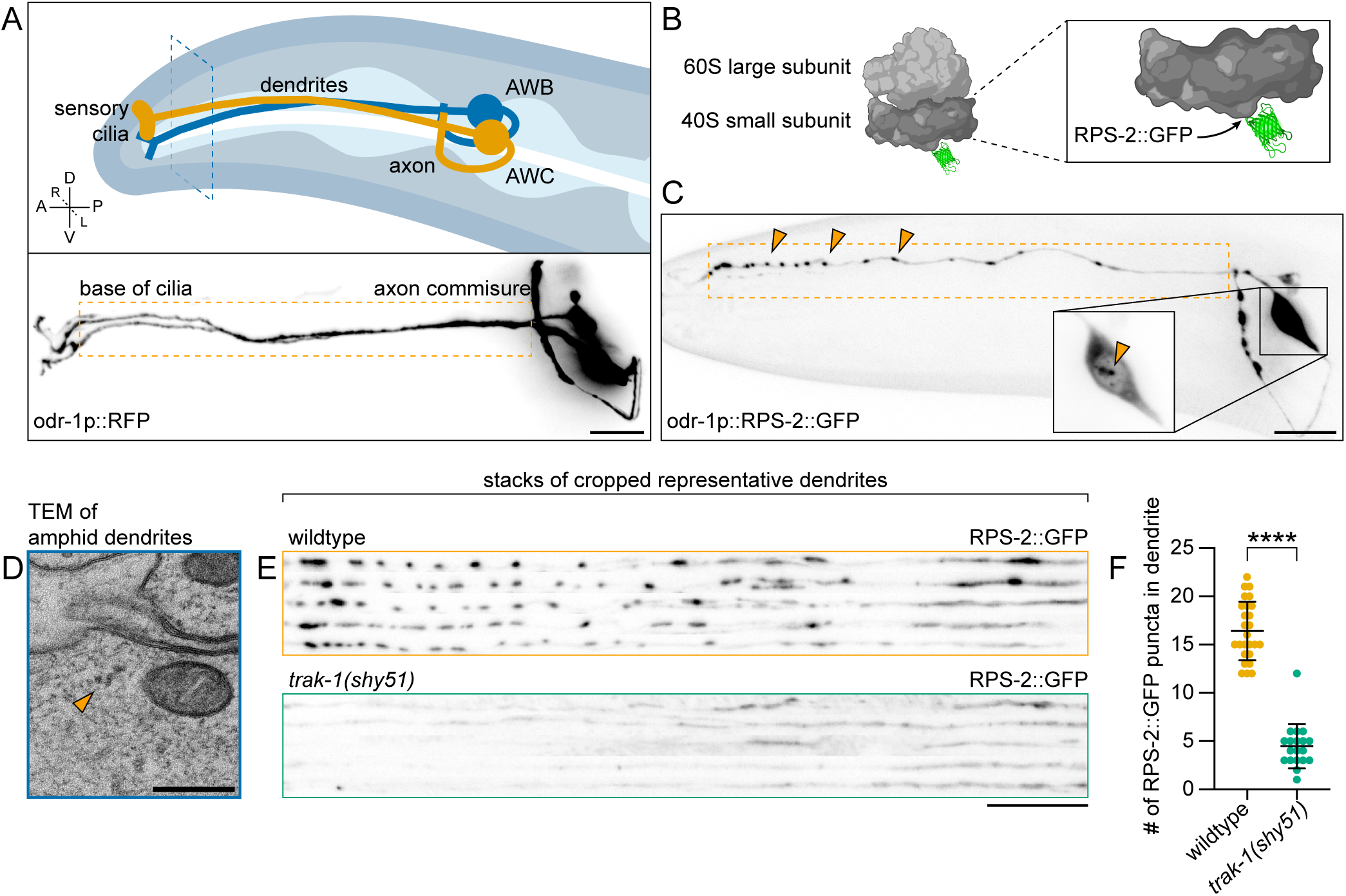
Ribosomes are stereotypically positioned in the dendrites of AWB and AWC. **(A)** Top: schematic of the chemosensory neurons AWB and AWC. Dashed blue lines indicate region presented in (D). Compass rose: D = dorsal, V = ventral, A = anterior, P = posterior, R = right, L = left. Bottom: cytosolic RFP expression in AWB and AWC driven by the *odr-1p* promoter. Dashed orange lines indicate the dendritic region presented throughout this study. Scale bar, 10 µm. **(B)** Schematic of RPS-2::GFP incorporation into the ribosome. **(C)** Representative confocal image (maximum intensity z-projection) of RPS-2::GFP localization in AWB and AWC neurons at larval stage L4. Dashed orange lines mark the dendrite, and orange arrowheads indicate dendritic RPS-2 puncta. Inset shows cell body, and orange arrowhead indicates nucleoli. Image is linearly adjusted for brightness and contrast and grayscale inverted. Scale bar, 10 µm. **(D)** Transmission electron micrograph of an amphid dendrite. Orange arrowhead indicates a polysome. Scale bar, 200 nm. **(E)** Representative images of RPS-2::GFP expression in wildtype animals compared to *trak-1(shy51)*. For each genotype, a stack of five representative dendrites is shown. Dendrites are straightened and cropped to the first 80 µm of the dendrite beginning from the base of the cilium. Scale bar, 10 µm. **(F)** Quantification of (E). Number of RPS-2::GFP puncta per dendrite in wildtype versus *trak-1(shy51)* mutants. Student’s t-test; n = 26 and 19 animals. Bars indicate mean and SD. ∗∗∗∗p < 0.0001

In the dendrites, RPS-2::GFP localizes to punctate signals along the length of the process (Fig. 1 C). Interestingly, dendritic RPS-2::GFP puncta are stereotypically positioned (Fig. 1 E shows an alignment of dendrites from several animals), suggesting the existence of mechanisms that regulate their localization. RPS-2::GFP is excluded from the sensory cilia, consistent with the absence of ribosomes in cilia visualized by electron microscopy (Doroquez et al., 2014). We could also detect RPS-2::GFP signal in the axon; however, as the axons of AWB and AWC receive postsynaptic inputs from other neurons (White et al., 1986), they are not strictly axonal and were not focused on here.

To validate the presence of dendritic ribosomes using an orthogonal approach, we performed electron microscopy in the distal region of the amphid dendrites and found that ribosomes are indeed present in this region (Fig. 1 D). These findings are consistent with a previous study of the endoplasmic reticulum in *C. elegans* neurons (Rolls et al., 2002). We also examined a previously-published electron microscopy dataset from an L1 stage larval worm (Witvliet et al., 2021). Consistent with our data, dendritic ribosomes are abundant in this dataset (Fig. S1 A). Taken altogether, we conclude that RPS-2::GFP recapitulates the expected distribution of ribosomes in AWB and AWC. Furthermore, dendritic ribosomes are abundant and localize to stereotypic puncta along the length of the dendrite.

### An unbiased genetic screen implicates a role for mitochondria in dendritic ribosome distribution

To identify regulators of ribosome transport and positioning in the dendrites of AWB and AWC, we performed an unbiased, forward genetic screen on worms expressing RPS-2::GFP to identify mutations that eliminate or mislocalize dendritic ribosome puncta. After screening 5,000 haploid genomes, we isolated and mapped (see Methods) two novel mutant alleles defective for punctate ribosome localization. First, we isolated *shy51*, a G478E missense allele of *trak-1*/TRAK1 (Fig. 1 E). In *trak-1(shy51)* mutant worms, the number of RPS-2::GFP puncta in the dendrite is strongly reduced (Fig. 1 F). We also isolated *shy54*, an A228T missense allele of *drp-1*/DNM1L (Fig. S1 B). In *drp-1(shy54)* mutant worms, RPS-2 signal becomes continuous along the proximal region of the dendrite and is specifically excluded from the distal dendrite (Fig. S1 C). Cell-specific expression of *DRP-1* in AWB and AWC could rescue this ribosome distribution defect, indicating that DRP-1 functions cell-autonomously (not shown).

TRAK-1 is well-known for its role in mitochondrial motility (Stowers et al., 2002). Furthermore, DRP-1 is a dynamin-like enzyme necessary for mitochondrial fission (Smirnova et al., 1998; Labrousse et al., 1999; Smirnova et al., 2001); mitochondria in *drp-1* mutants undergo drastic changes in morphology due to excessive fusion events (Labrousse et al., 1999; Byrne et al., 2019). The independent isolation of two genes involved in mitochondria transport and morphology suggested a role for mitochondria in dendritic ribosome positioning in AWB and AWC.

### Ribosomes colocalize with mitochondria in the dendrite

To investigate the potential roles played by mitochondria in dendritic ribosome positioning, we co-expressed reporters for ribosomes and mitochondria under the *odr-1p* promoter. To visualize the mitochondrial matrix, we fused a codon-optimized sequence encoding the 24aa mitochondrial localization sequence of chicken mitochondrial aspartate aminotransferase (Jaussi et al., 1985) and a short linker peptide to the N-terminus of TagRFP-T (hereafter referred to as “mitoRFP”). RPS-2::GFP and mitoRFP exhibit a strikingly high degree of colocalization in the dendrite (Fig. 2 A) but not in the soma (Fig. 2 B), which is reflected by linescan comparisons of normalized fluorescence (Fig. 2, C and D) and the Pearson’s Correlation Coefficient (Fig. 2 E). To validate that ribosomes localize to dendritic mitochondria using an orthogonal approach, we examined our electron microscopy data and noted examples of polysomes in close proximity to mitochondria (Fig. 1 D). To extend these findings further, we again examined cross sections through the amphid dendrites in a previously-published electron microscopy dataset (Witvliet et al., 2021). Mitochondria with closely associated ribosomes are common in the amphid dendrites (Fig. 2 F). For such mitochondria, ribosomes and polysomes are frequently observed to be located at an average of ~20 nm (SD = 14 nm) away (Fig. 2 G).

**Figure 2.**
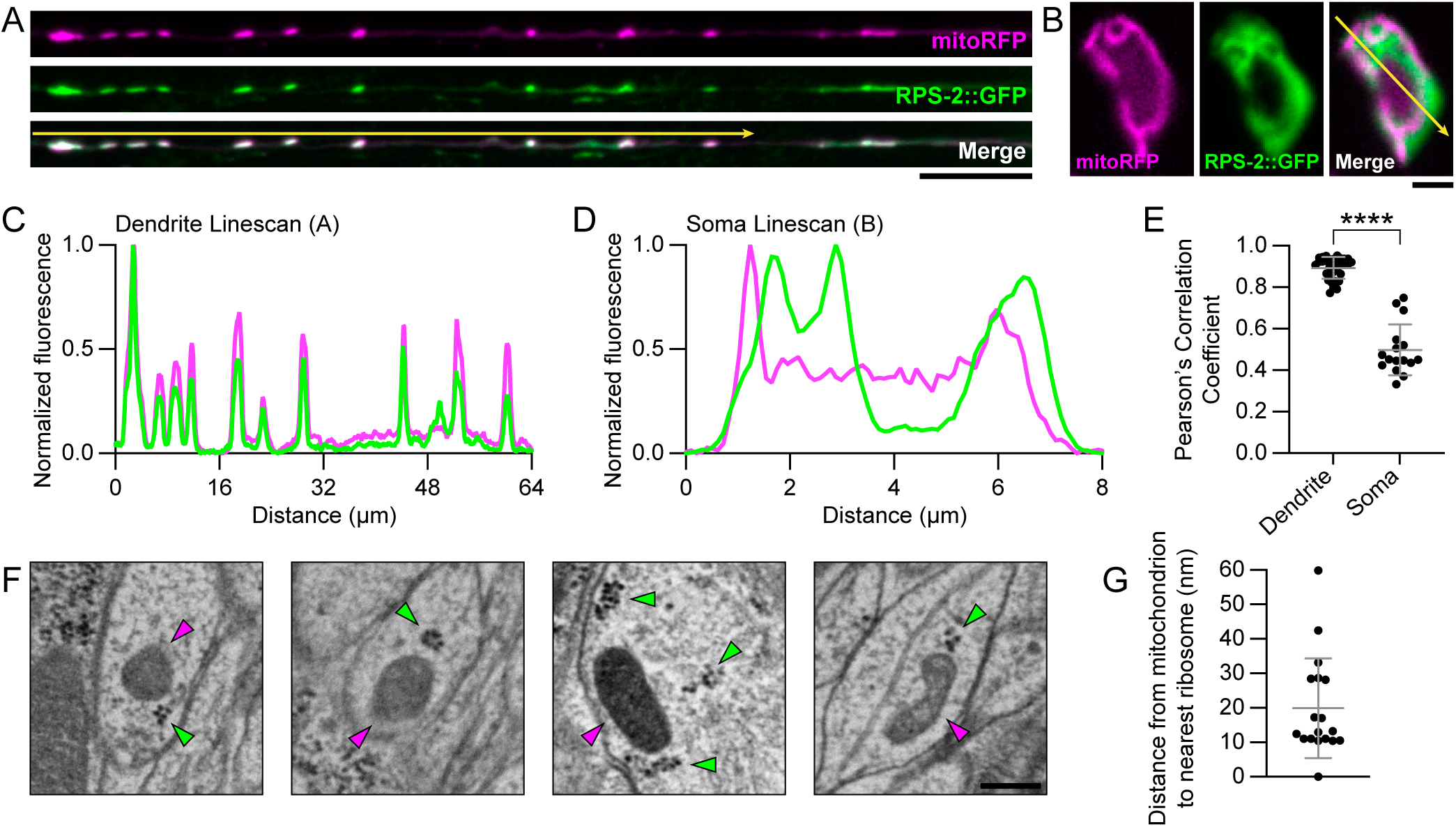
Ribosomes colocalize with mitochondria in the dendrite. **(A)** Confocal images of mitoRFP and RPS-2::GFP colocalization in the dendrite. Yellow arrow indicates region of linescan in (C) The linescan was measured exactly 1 µm beneath the yellow arrow. Scale bar, 10 µm. **(B)** Confocal images of mitoRFP and RPS-2::GFP localization in a single focal plane of the AWB cell body. Yellow arrow indicates region of linescan in (D). Scale bar, 2 µm. **(C-D)** Linescans of normalized fluorescence intensity across the dendrite (C) or AWB cell body (D) indicated by yellow arrows in (A) and (B). **(E)** Pearson’s Correlation Coefficient quantifying (A) and (B). Student’s t-test; n = 32 and 16 measurements. Soma measurements taken from subset of animals with dendrite measurements. Bars indicate mean and SD. **(F)** Representative scanning electron microscopy images of mitochondria with associated polysomes in oblique cross sections through the amphid dendrites. Magenta arrowheads indicate mitochondria; green arrowheads indicate polysomes. Data from Witvliet et al. (2021). Scale bar, 200 nm. **(G)** Quantification of (F). n = 18 mitochondria from 1 animal; bars indicate mean and SD. ∗∗∗∗p < 0.0001

### Dendritic ribosome puncta depend on mitochondria

We asked if the steady-state localization of dendritic ribosomes depends on mitochondria transport. To address this question, we assessed the effects of mutant alleles for components of the mitochondrial transport machinery on the localization of mitochondria and ribosomes in the dendrite. As the minus-ends of microtubules are uniformly oriented away from the cell body in the dendrites of AWB and AWC (Maniar et al., 2012), microtubule-based transport of mitochondria towards the tip of the dendrite is likely driven by dynein (schematized in Fig. 3 A). To disrupt dynein motor activity, we first tested a nonsense allele of the dynein light intermediate chain *dli-1*/DYNC1LI2 (Zhao et al., 2021). This allele causes a significant reduction in the number of dendritic mitochondria, and in contrast to wildtype animals (Fig. 3 B), mitochondria in *dli-1* are specifically excluded from the distal region of the dendrite (Fig. 3, C and I). This reduction and mislocalization of mitochondria is accompanied by a concomitant mislocalization and decrease in the number of dendritic RPS-2 puncta as well (Fig. 3, C and J).

**Figure 3.**
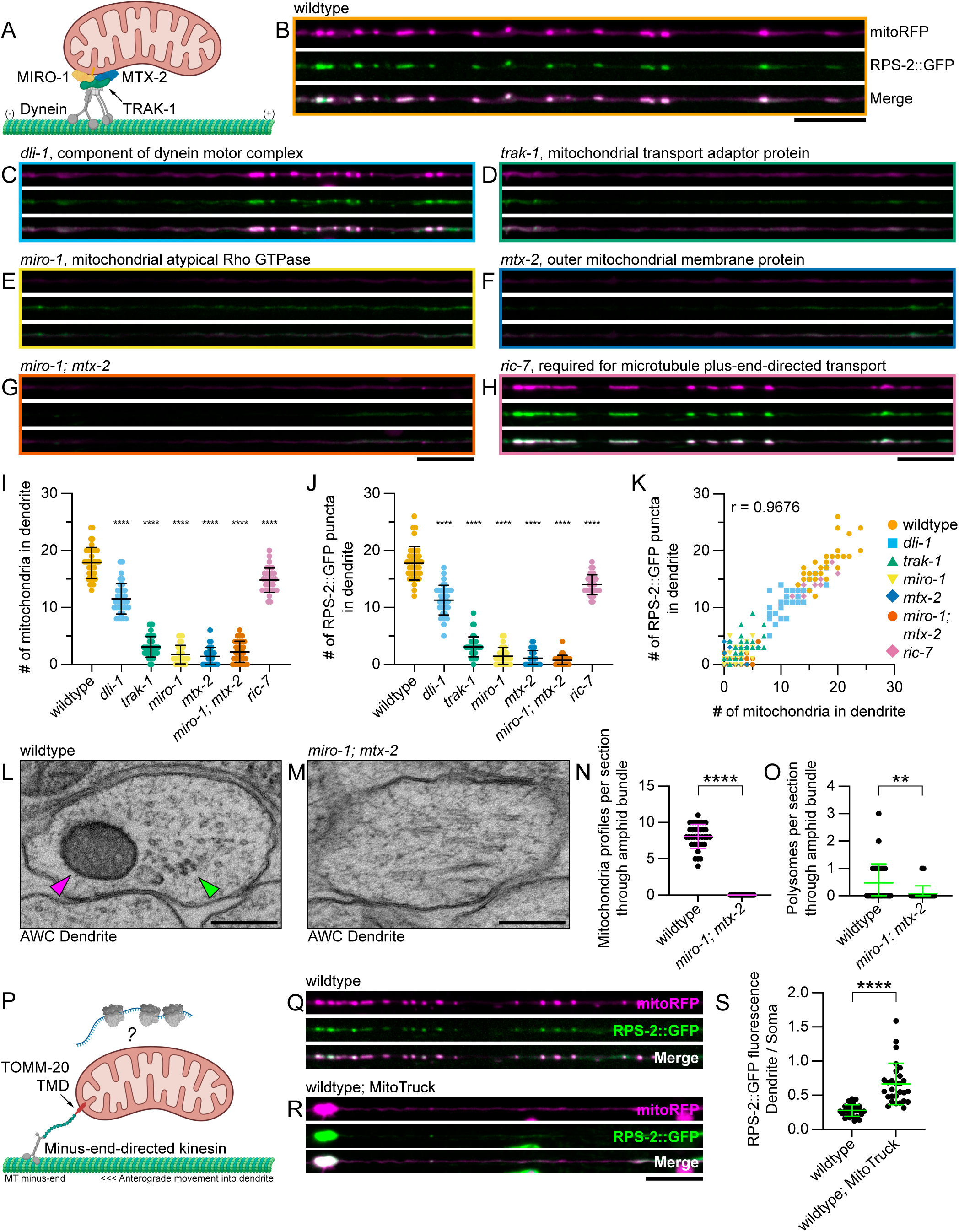
Mitochondria are necessary and instructive for dendritic ribosome positioning. **(A)** Model of adaptor complex for microtubule minus-end-directed mitochondria transport in *C. elegans*, as proposed by Zhao et al. (2021). **(B-H)** Representative confocal images demonstrating localization of mitoRFP and RPS-2::GFP in the dendrites of wildtype (B), *dli-1* (C), *trak-1* (D), *miro-1* (E), *mtx-2* (F), *miro-1; mtx-2* (G), and *ric-7* (H) worms. Scale bar, 10 µm. **(I and J)** Quantification of (B-H). n = 29-39 animals. One-way ANOVA with Dunnett’s multiple comparisons test. Bars indicate mean and SD. Comparison of dendritic mitochondria counts (I) or RPS-2 puncta (J) in the dendrite in each genetic background tested. **(K)** Same data from (I) and (J) plotted against each other to compare the numbers of dendritic mitochondria and RPS-2 puncta. Pearson’s Correlation Coefficient; r(230)=0.9676, p<0.0001. **(L and M)** Representative transmission electron microscopy images of cross-sections through the AWC dendrite in a wildtype (L) and *miro-1; mtx-2* double mutant (M) worm. Magenta arrowheads indicate mitochondria; green arrowheads indicate polysomes. Scale bar, 200 nm. **(N and O)** Quantification of (L) and (M). The number of mitochondria profiles (N) or polysomes (O) observed per 55 nm cross-section through the left amphid bundle (comprised of twelve dendrites). Nonparametric Mann-Whitney test, n = 36 serial sections at 55 nm thickness (~2 µm total depth). Bars indicate mean and SD. **(P)** Schematic of chimeric MitoTruck construct. **(Q and R)** Representative confocal images of dendritic mitoRFP and RPS-2::GFP localization in indicated genotypes. **(S)** Quantification of (Q) and (R). n = 14-30 animals. Student’s t-test; n = 30 and 28 animals. Bars indicate mean and SD. ∗∗∗∗p < 0.0001; ∗∗p < 0.01

Previous studies have identified *trak-1*/TRAK1 (Stowers et al., 2002), *miro-1*/MIRO-1 (Fransson et al., 2003, 2006; Guo et al., 2005), and *mtx-2*/Metaxin-2 (Zhao et al., 2021) as regulators of dynein-mediated mitochondria transport in dendrites (schematized in Fig. 3 A). Consistent with these studies, null alleles of *trak-1(wy50182)* (Zhao et al., 2021), *miro-1(wp88)* (Wu et al., 2024), and *mtx-2(wy50266)* (Zhao et al., 2021) all strongly reduce the number of mitochondria in the dendrite (Fig. 3 D-F, I). Notably, in each of these independent mutant backgrounds, the loss of mitochondria is accompanied by a loss of dendritic RPS-2 puncta as well (Fig. 3 D-F, J). We also tested a *miro-1; mtx-2* double mutant, which exhibits a phenotype similar to both single mutants (Fig. 3, G, I, and J).

As a negative control, we selectively disrupted mitochondria transport towards the plus-ends of microtubules using a nonsense allele of *ric-7(n2657)*, a gene required for the kinesin-1-mediated transport of mitochondria towards microtubule plus-ends in *C. elegans* (Rawson et al., 2014; Wu et al., 2024). As expected, loss of *ric-7* does not eliminate dendritic mitochondria (Fig. 3, H and I). We did observe a slight decrease in the number of dendritic mitochondria, and the average length of mitochondria in the distal dendrite was increased (not shown). This may reflect impaired retrograde transport back to the cell body and an overall increase in dendritic mitochondrial mass, leading to excessive fusion events. Notably, in this transport mutant background where dendritic mitochondria are retained, dendritic RPS-2 puncta also remain (Fig. 3, H and J).

Taken altogether, these data strongly suggest that dendritic ribosome localization is dependent upon mitochondria in this system. Indeed, the numbers of dendritic RPS-2 puncta and mitochondria observed by fluorescence microscopy across all genotypes are highly correlated, Pearson’s r(230)=0.9676, p<0.0001 (Fig. 3 K). To extend these findings further with an orthogonal approach, we performed electron microscopy to compare ribosome and mitochondria localization in the amphid dendrites between a wildtype and *miro-1; mtx-2* double mutant worm (Fig. 3 L-M). In L4 larval stage worms, we examined a depth of ~2 µm in serial cross sections through the left amphid bundle (36 sections of 55 nm thickness each), beginning shortly past the base of the sensory cilia and continuing posteriorly through the distal amphid, as our fluorescence imaging had demonstrated this distal dendrite region to be abundant with mitochondria (Fig. 3 B). We focused our analysis on polysomes, as careful examination of consecutive cross sections revealed that granular particles resembling monosomes are often continuous across sections, suggesting that they are in fact filamentous structures rather than monosomes. As expected, the number of observed dendritic mitochondria is strikingly reduced in the *miro-1; mtx-2* double mutant (Fig. 3 N). We also observed a concomitant strong decrease in the number of dendritic polysomes (Fig. 3 O). These electron microscopy data are consistent with a model in which mitochondria play a role in dendritic ribosome localization.

### Mitochondria are sufficient to instruct ribosome localization in the dendrite

Our loss-of-function mutant analyses provide strong genetic evidence that dendritic ribosome localization is dependent on mitochondria. To extend these findings further, we asked if mitochondria can also be instructive for dendritic ribosome localization in a gain-of-function context. Inspired by chimeric motor protein fusions previously used to re-localize mitochondria and other organelles in neurons (Rawson et al., 2014; Farías et al., 2019; Jia and Sieburth, 2021), we designed the artificial tethering construct TOMM-20TMD::Linker::KLP-16(T128-N587), hereafter referred to as “MitoTruck” (Fig. 3 P). MitoTruck fuses the motor domain of the minus-end-directed kinesin KLP-16/KIFC1/kinesin-14 to the N-terminal 55aa transmembrane domain of TOMM-20/TOM20 in order to ectopically drag mitochondria towards the minus-end of microtubules.

When expressed under the *odr-1p* promoter, MitoTruck drags mitochondria to the distal end of the dendrite (Fig. 3 R), causing a large swelling of the dendrite tip. While a few mitochondria occasionally remain in the medial dendrite, the majority of mitochondria along the length of the dendrite are lost. Notably, MitoTruck expression also leads to a large accumulation of RPS-2::GFP signal at the tip of the dendrite (Fig. 3 R), resulting in a significant increase in dendritic RPS-2::GFP signal compared to the cell body (Fig. 3 S). These results demonstrate that mitochondria are instructive for dendritic ribosome localization. We conclude that mitochondria are both necessary and instructive for dendritic ribosome positioning in this system.

### Ribosomes co-transport with a distinct population of mitochondria in the dendrite

Having established the steady-state relationship between ribosomes and mitochondria, we next asked if ribosomes can co-transport with mitochondria. Timelapse imaging reveals that ribosomes do indeed co-transport with mitochondria in both the proximal (Fig. 4 A) and distal (Fig. S2 A) regions of the dendrite. Of all detectable RPS-2 movements, 74% have associated mitochondria (Fig. S2 B), suggesting that mitochondrial motility can explain a majority of ribosome transport events in this system.

**Figure 4.**
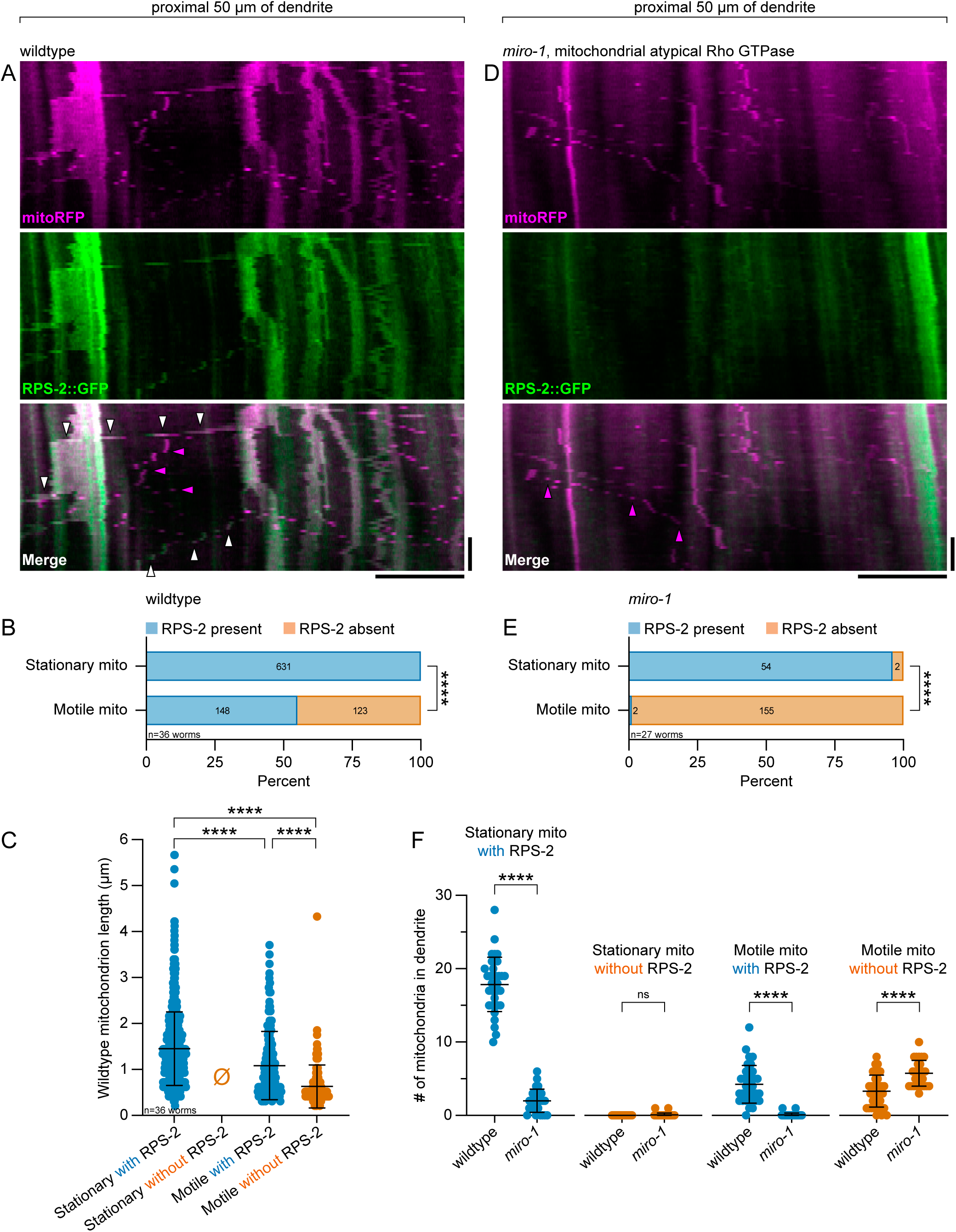
*miro-1* dependent co-transport of ribosomes with mitochondria. **(A)** Representative kymographs of mitoRFP and RPS-2::GFP movements in the proximal dendrite, generated from a 10 min movie with a 10 sec framerate. White arrowheads indicate examples of co-transport between RPS-2::GFP and mitoRFP. Magenta arrowheads indicate example of a motile mitochondrion lacking associated RPS-2::GFP. Horizontal scale bar, 10 µm; vertical scale bar, 2 min. **(B-C)** Quantification of (A), n=36 animals.. **(B)** Fisher’s exact test comparing the relationship between motility and RPS-2 association amongst mitochondria (two-tailed p <0.0001). **(C)** Quantification of wildtype mitochondrion length grouped by motility and presence of RPS-2. n = 631, null, 146, and 123 mitochondria from 36 animals. One-way ANOVA with Tukey’s post-hoc test. Bars indicate mean and SD. **(D)** Representative kymographs of dendritic mitoRFP and RPS-2::GFP movement in a *miro-1* mutant background. Magenta arrowheads indicate example of a motile mitochondrion lacking associated RPS-2::GFP. Horizontal scale bar, 10 µm; vertical scale bar, 2 min. **(E)** Quantification of (D). Fisher’s exact test as in (B). **(F)** Quantification of the number of mitochondria per dendrite in wildtype versus *miro-1* mutants, grouped by motility and presence of RPS-2. Student’s t-tests; n = 35 animals for wildtype and 27 animals for *miro-1*. ns = not significant; ∗∗∗∗p < 0.0001

Our timelapse imaging also enabled the identification of many small, motile mitochondria that are otherwise difficult to detect above background signal in steady-state images (Fig. 4 A). We noticed the presence of motile mitochondria lacking RPS-2, raising the possibility that RPS-2 might only associate with a subset of dendritic mitochondria. Quantification reveals that only 55% of all motile mitochondria have associated RPS-2 (Fig. 4 B). The majority of motile mitochondria were observed in the proximal dendrite (Fig. 4 A); in contrast, most mitochondria in the distal dendrite were stationary and only rarely observed moving (Fig. S2 A). Strikingly, 100% of these stationary mitochondria have associated RPS-2 (Fig. S2 A); in 6 hours of video from 36 worms we did not observe a stationary mitochondrion without it (Fig. 4 B).

In addition to their marked difference in RPS-2 association, we also noted a difference in mitochondrion size between stationary and motile mitochondria. Measuring mitochondrion length reveals that stationary mitochondria are larger than motile mitochondria, and that amongst motile mitochondria, those with associated RPS-2 tend to be larger than the mitochondria lacking any associated RPS-2 (Fig. 4 C). Altogether, these data suggest the presence of multiple distinct populations of mitochondria in the dendrites of AWB and AWC: 1) a population of large, stationary mitochondria in the distal dendrite with associated ribosomes, 2) a population of smaller motile mitochondria also bearing ribosomes, observed throughout the dendrite, and 3) a population comprised of the smallest mitochondria, most commonly observed in the proximal and medial dendrite, which are highly motile and lack associated ribosomes. Thus, the majority of ribosome movement events in this system are mediated by a subset of motile mitochondria.

### Co-transport between ribosomes and mitochondria requires *miro-1*

We next performed timelapse imaging in *miro-1* mutants (Fig. 4 D and Fig. S2 C), as our steady-state imaging had revealed a strong loss of mitochondria and RPS-2 puncta in this background (Fig. 3 E, I, and J). In *miro-1* mutants, the large, stationary mitochondria and their associated ribosomes are mostly lost from the distal dendrite (Fig. S2 C and Fig. 4, E and F). In contrast, the total number of motile mitochondria is only modestly reduced from an average of ~8 motile mitochondria per dendrite in wildtype animals to an average of ~6 per dendrite in *miro-1* mutants (compare Fig. 4 B and E). Strikingly, however, nearly all of the motile mitochondria in *miro-1* mutants lack associated RPS-2 (Fig. 4 E), suggesting that *miro-1* is required for the association of ribosomes with motile mitochondria. Consistent with this idea, while the average length of each mitochondrion class is comparable to wildtype (Fig. S2 E), the relative abundance of each mitochondrial population is significantly altered by loss of *miro-1*. Whereas RPS-2-positive motile mitochondria are decreased, RPS-2-negative motile mitochondria became more abundant (Fig. 4 F).

Finally, we noted that while the abundance of motile RPS-2 puncta in *miro-1* mutants is considerably decreased, most of the remaining RPS-2 movements lack associated mitochondria (Fig. S2 D). Such movements were also visible in wildtype animals (Fig. S2 B), demonstrating the existence of a small population of mitochondria-independent motile RPS-2 puncta.

### RAB-7 colocalizes with RPS-2 and mitochondria in the dendrite

Previous studies have implicated several membrane-bound organelles as drivers of mRNA and ribosome transport through a mechanism of “organelle hitchhiking” (Higuchi et al., 2014; Cioni et al., 2019; Liao et al., 2019; Harbauer et al., 2022; Cohen et al., 2022); therefore, we sought to clarify the roles played by mitochondria, endosomes, and lysosomes in dendritic ribosome localization by visualizing markers for each organelle in the dendrite. To enable three-color imaging, we cloned markers with a codon-optimized sequence for mTagBFP2 (hereafter referred to as “BFP”). BFP::RAB-5, a reporter for early endosomes, is most often present in the proximal dendrite (Fig. 5 A). We observed an average of ~7 early endosomes per dendrite (Fig. 5 B), which is notably less abundant than dendritic RPS-2::GFP puncta (~18 per dendrite, Fig. 3 J). Furthermore, over half of the worms imaged lacked RAB-5 puncta in the distal region of the dendrite (Fig. S3 A) where RPS-2 puncta are abundant. These results suggest that early endosomes are unlikely to mediate ribosome transport in these dendrites. Consistent with previous reports (Cioni et al., 2019; Schuhmacher et al., 2023), the RAB-5 puncta that were present in the dendrite displayed an 80% overlap with RPS-2 (Fig. 5 B).

**Figure 5.**
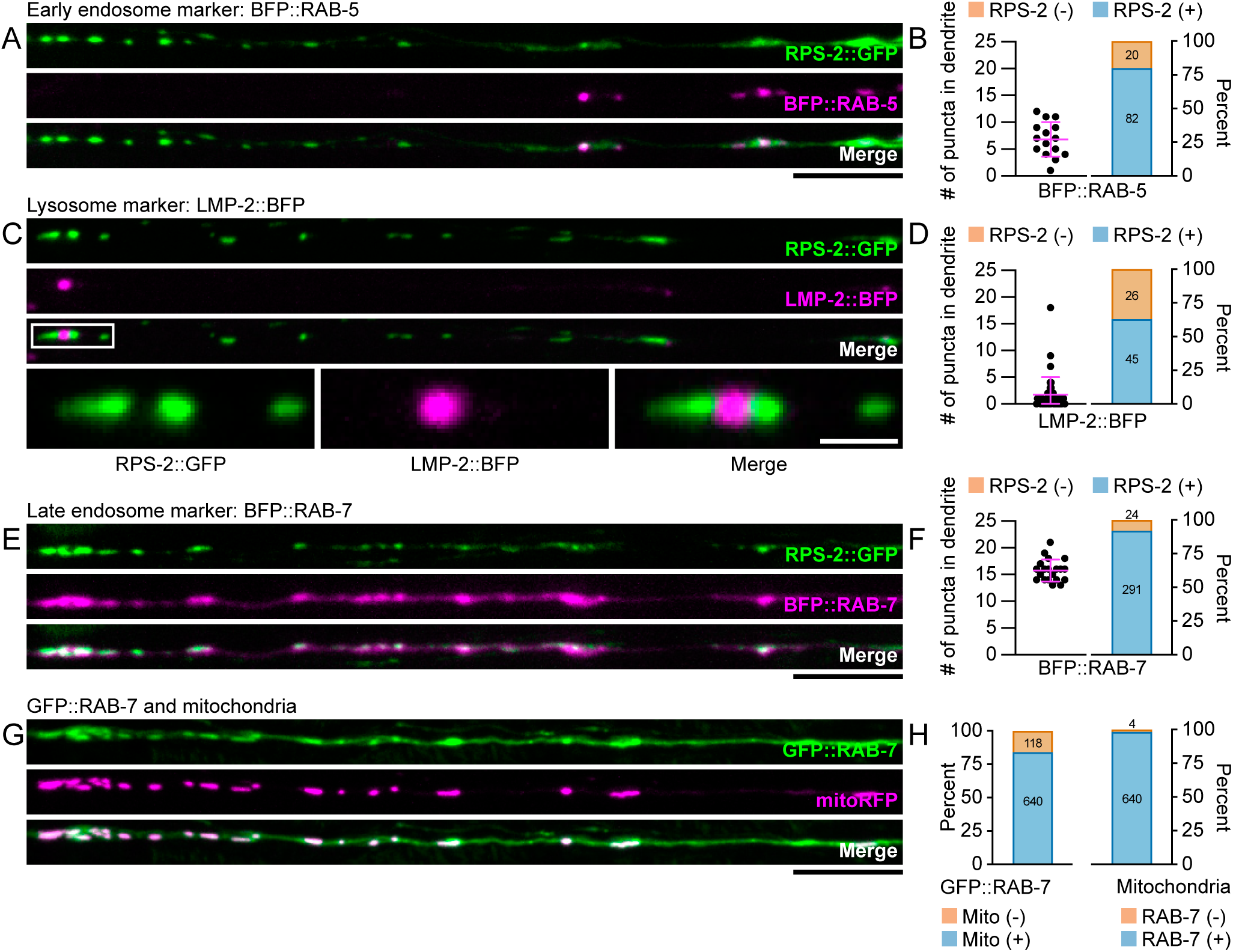
RAB-7 colocalizes with RPS-2 and mitochondria in the dendrite. **(A)** Confocal images of RPS-2::GFP and BFP::RAB-5 localization in the dendrite. Scale bar, 10 µm. **(B)** Quantification of (A). Number of BFP::RAB-5 puncta in the dendrite, and percentage of BFP::RAB-5 puncta with “(+)” and without “(-)” associated RPS-2::GFP signal. n = 15 animals. **(C)** Confocal images of RPS-2::GFP and LMP-2::BFP localization in the dendrite. Above: scale of images is the same as (A). White box indicates region of inset. Below, inset: high-magnification view of RPS-2::GFP and LMP-2::BFP in the distal dendrite. White scale bar, 2 µm. **(D)** Quantification of (C). Number of LMP-2::BFP puncta in the dendrite, and percentage of LMP-2::BFP puncta with and without associated RPS-2::GFP signal. n = 40 animals. **(E)** Confocal images of RPS-2::GFP and BFP::RAB-7 localization in the dendrite. Scale bar, 10 µm. **(F)** Quantification of (E). Number of BFP::RAB-7 puncta in the dendrite, and percentage of BFP::RAB-7 puncta with and without associated RPS-2::GFP signal. n = 20 animals. **(G)** Confocal images of GFP::RAB-7 and mitoRFP localization in the dendrite. Scale bar, 10 µm. **(H)** Quantification of (G). Percentage of GFP::RAB-7 puncta with and without associated mitochondria, and percentage of mitochondria with and without associated GFP::RAB-7 signal. n = 31 animals.

We also imaged LMP-2::BFP, the closest worm homolog of the lysosome reporter LAMP1 (Fig. 5 C). We found that dendritic lysosomes are rare in this system, with an average of ~2 LMP-2 puncta per dendrite (Fig. 5 D), and with 40% of worms lacking detectable LMP-2::BFP signal in the dendrite (Fig. S3 B). In the worms that do exhibit dendritic LMP-2 signal, 63% of LMP-2 vesicles overlap with RPS-2 (Fig. 5 D). LMP-2 vesicles not overlapping with RPS-2 could occasionally be observed flanking RPS-2 puncta (Fig. 5 C). These data suggest that lysosomes are also unlikely to play a major role in determining ribosome localization in this system.

In contrast, the late endosome reporter BFP::RAB-7 exhibits a high degree of colocalization with RPS-2::GFP in the dendrite (Fig. 5 E). Unlike RAB-5 and LMP-2, which localize to discrete puncta, we observed BFP::RAB-7 signals that are both diffuse across the length of the dendrite and enriched at puncta localizing to RPS-2::GFP (Fig. 5 E). Considering that the majority of these BFP::RAB-7 puncta overlap with RPS-2::GFP signal (Fig. 5 F) and that the average number of BFP::RAB-7 puncta per dendrite corresponds well to the average number of dendritic mitochondria (Fig. 5 F and Fig. 3 I), we asked if RAB-7 also colocalizes with mitochondria in the dendrite. Similar to BFP::RAB-7, we observed diffuse GFP::RAB-7 signal throughout the dendrite (Fig. 5 G). We also observed regions where GFP::RAB-7 signal enriched into puncta, which colocalized strongly with mitochondria (Fig. 5, G and H). While most dendritic mitochondria are enriched with GFP::RAB-7 signal, 16% of GFP::RAB-7 puncta lack associated mitochondria (Fig. 5 H), suggesting that RAB-7 might localize to multiple structures in the dendrite.

To test this idea further, we performed timelapse imaging of GFP::RAB-7 and mitoRFP (Fig. S3 C). We found that GFP::RAB-7 can co-transport with mitochondria, and the majority of both stationary and motile mitochondria possess associated RAB-7 signal (Fig. S3 D). Similarly, the majority of stationary RAB-7 signals are associated with mitochondria; however, a large fraction of motile RAB-7 signals lack associated mitochondria (Fig. S3 E). These data demonstrate that while RAB-7 strongly colocalizes with mitochondria in the dendrite, it is also present on highly motile vesicles that are transported independently of mitochondria. We conclude that while RAB-5-positive and LMP-2-positive vesicles are not located closely enough to RPS-2 puncta to play a major role in dendritic ribosome localization, RAB-7 signal is closely associated with both RPS-2 and mitochondria in the dendrite.

### Ribosome localization to mitochondria is independent of RAB-7 enrichment

Because previous studies have reported roles for late endosomes and lysosomes in the positioning of mRNAs and ribosomes near mitochondria in axons (Cioni et al., 2019; Liao et al., 2019; De Pace et al., 2024), we considered the possibility that RAB-7-positive vesicles might position ribosomes near mitochondria in this system. In this model, mitochondria would play an indirect role in ribosome localization by instructing the positioning of RAB-7-positive vesicles in the dendrite. To test this model, we sought to genetically dissociate RAB-7 signal from mitochondria in order to determine if RPS-2 localization at mitochondria is dependent on the presence of RAB-7. First, we attempted to disrupt RAB-7 colocalization by perturbing mitochondria positioning. In *miro-1* deletion mutants, BFP::RAB-7 puncta are lost from the dendrite in addition to mitochondria (Fig. 6, A-C), demonstrating that mitochondria are required for the localization of RAB-7 in the dendrite. We then tested RAB-7 localization in a MitoTruck background. Ectopic re-localization of mitochondria to the dendrite tip also re-localizes BFP::RAB-7 (Fig. 6, D and E). These data highlight the close association between RAB-7 and mitochondria in the dendrite.

**Figure 6.**
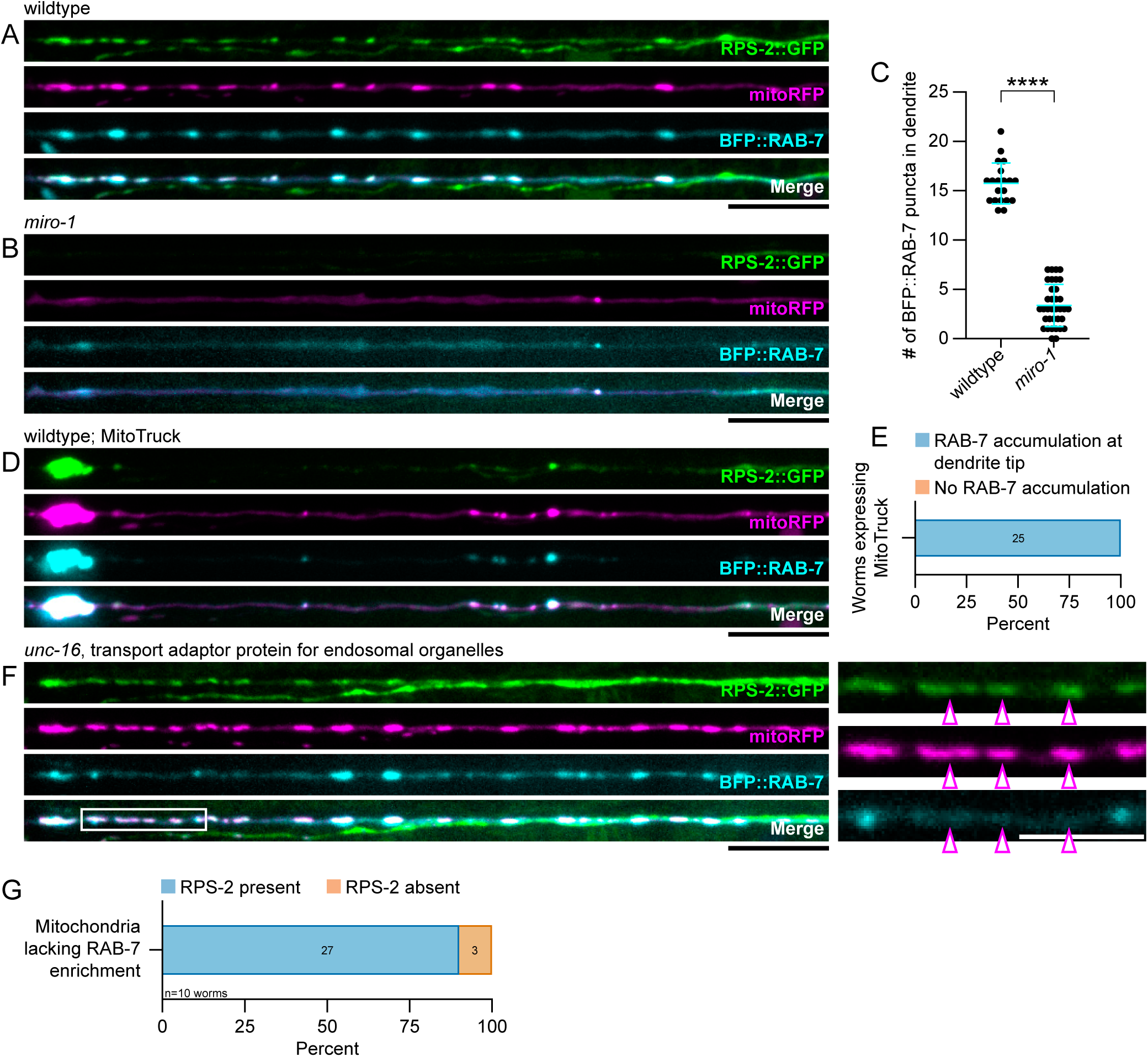
Ribosome localization to mitochondria is independent of RAB-7 enrichment. **(A and B)** Confocal images of dendritic RPS-2::GFP, mitoRFP, and BFP::RAB-7 localization in wildtype (A) and *miro-1* (B) genetic backgrounds. **(C)** Quantification of (A) and (B). Number of BFP::RAB-7 puncta per dendrite in wildtype versus *miro-1* mutants. Student’s t-test; n = 20 and 35 animals. Bars indicate mean and SD. The puncta counts for wildtype animals are the same as those displayed in Figure 5 F. **(D)** As in (A), with additional MitoTruck expression. **(E)** Quantification of (D). Percentage of MitoTruck-expressing worms with and without an accumulation of BFP::RAB-7 at the tip of the dendrite. n = 25 animals. **(F)** Confocal images of RPS-2::GFP, mitoRFP and BFP::RAB-7 in the *unc-16* mutant background. White box indicates region of inset. Right, inset: high-magnification view of all three markers in the distal dendrite. Hollow magenta arrowheads indicate mitochondria with associated RPS-2::GFP that lack enrichment of BFP::RAB-7 signal. White scale bar, 5 µm. (G) Quantification of (F). Of mitochondria in *unc-16* mutants that lack BFP::RAB-7 enrichment, percentage with and without associated RPS-2. n = 10 animals. ∗∗∗∗p < 0.0001. Black scale bars, 10 µm.

In wildtype animals, we occasionally observed instances of mitochondria lacking GFP::RAB-7 enrichment (Fig. 5 H). We reasoned that mutations disrupting RAB-7 entry into the dendrite might further dissociate RAB-7 from mitochondria. Previous studies have identified UNC-16/JIP3 as a scaffold protein that can serve as an adaptor between endosomal organelles and dynein (Cavalli et al., 2005; Arimoto et al., 2011). Speculating that loss of UNC-16/JIP3 might reduce RAB-7 signal the dendrite, we tested a full deletion allele of *unc-16(prt183)* (Celestino et al., 2022), the sole worm homolog of mammalian JIP3/MAPK8IP3 and JIP4/SPAG9. Dendritic BFP::RAB-7 signal is not eliminated in *unc-16* mutants (Fig. 6 F); however, we observed a low-penetrance dissociation phenotype between RAB-7 and mitochondria. In a subset of *unc-16* worms, mitochondria occasionally lack an enrichment of BFP::RAB-7 signal (Fig. 6 F). This dissociation is visible in a strain expressing GFP::RAB-7 as well (Fig. S4, A and B), demonstrating the effect is not due to low visibility of the marker.

We took advantage of the dissociation phenotype in *unc-16* mutants to ask if RPS-2 can localize to mitochondria in the absence of RAB-7 enrichment. We observed that mitochondria lacking BFP::RAB-7 enrichment can still possess associated RPS-2::GFP (Fig. 6 F). After blindly scoring such RAB-7-negative mitochondria for the presence or absence of RPS-2, we found that 90% of mitochondria lacking BFP::RAB-7 enrichment still possess associated RPS-2::GFP signal (Fig. 6 G). From these data, we conclude that RPS-2 localization to mitochondria does not depend on the presence of enriched RAB-7 signal; therefore, we consider it unlikely that mitochondria merely serve to position ribosome-bearing endolysosomal vesicles in the dendrite. Taken altogether, the colocalization and co-transport of mitochondria with ribosomes, as well as the requirement and sufficiency of mitochondria for instructing ribosome positioning, suggest that ribosomes hitchhike on mitochondria for their delivery to a sensory dendrite in *C. elegans*.

## DISCUSSION

How ribosomes are transported to dendrites *in vivo* is unclear. Here we report a key role for mitochondria in dendritic ribosome transport and positioning. An unbiased forward genetic screen for regulators of ribosome positioning yielded hits in genes that govern mitochondria transport and morphology. We find that ribosomes colocalize with mitochondria in the dendrites of sensory neurons, an observation supported by fluorescence and electron microscopy. Genetic loss of mitochondria from the dendrite results in a concomitant loss of ribosomes; furthermore, ectopic re-localization of mitochondria is sufficient to re-localize ribosomes, demonstrating that mitochondria are instructive for dendritic ribosome positioning. Finally, timelapse imaging reveals that 74% of all detectable ribosome movements in the dendrite are with mitochondria and that *miro-1* is important for this association. Taken altogether, these results support a model in which ribosomes hitchhike on mitochondria from neuronal soma to dendrites.

Previous studies that primarily focused on mRNA transport identified roles for early endosomes (Schuhmacher et al., 2023), late endosomes (Cioni et al., 2019), lysosomes (Liao et al., 2019; De Pace et al., 2024), and mitochondria (Harbauer et al., 2022; Cohen et al., 2022). We specifically compared the distribution of early endosomes and lysosomes to dendritic ribosomes *in vivo* and found that they were not as strongly colocalized as mitochondria, suggesting that these organelles do not instruct ribosome positioning. Furthermore, mutant alleles for worm homologs of annexin A11, a protein that tethers mRNP granules to lysosomes in axons (Liao et al., 2019), do not remove RPS-2::GFP signal from the dendrite (not shown). While RAB-7 colocalizes well with mitochondria and ribosomes in control animals, its dissociation from mitochondria in *unc-16/jip3* mutants does not disrupt the tight association between mitochondria and ribosomes, suggesting that it is not involved in ribosome distribution. We did observe that 26% of motile RPS-2 signals lack an associated mitochondrion, demonstrating the existence of mitochondrion-independent ribosome transport in the dendrite. For such events, mRNP granules (Knowles et al., 1996; Kanai et al., 2004; Elvira et al., 2006; Graber et al., 2013, 2017; Fukuda et al., 2020) and endolysosomal organelles could play an auxiliary role. These results highlight the existence of various strategies for cells to mobilize ribosomes, raising the intriguing possibility that specialized delivery methods are utilized in different cell types or for targeting to distinct subcellular locations.

How do ribosomes associate with mitochondria? In yeast, translating ribosomes bind to the outer mitochondrial membrane as their nascent peptide chain is translocated though the TOM complex (Kellems et al., 1974, 1975; Gold et al., 2017; Chang et al., 2024). In our electron micrographs and in a published dataset (Witvliet et al., 2021), we did not frequently observe polysomes directly contacting mitochondria; however, we note that these data cannot entirely rule out such a model as monosomes on the surface of mitochondria would not be reliably detected in our datasets (see Methods). Nevertheless, our observation that polysomes localize to an average of ~20 nm from the outer mitochondrial membrane raises the possibility that a long molecule such as mRNA could act as a tether.

Furthermore, our timelapse imaging experiments demonstrate that *miro-1* is genetically required for ribosomes to hitchhike on motile mitochondria in the dendrite. Hence, MIRO-1’s activity as an atypical Rho GTPase (Fransson et al., 2003, 2006) might enable it to recruit a downstream effector protein to the outer mitochondrial membrane to serve as an anchor for ribosome transport. A regulated association between mitochondria and ribosomes would also be consistent with our observation that not all motile mitochondria co-transport ribosomes. It is tempting to speculate that recruitment of an RNA-binding protein (RBP) to the outer mitochondrial membrane could serve to tether mRNA-bound ribosomes during transport. Several RBPs are known to localize to the surface of mitochondria (Béthune et al., 2019), and one was recently shown to specifically mediate the mitochondrial localization of neuronal *Pink1* mRNA in a translation-dependent manner (Harbauer et al., 2022). As mutations in worm homologs of SYNJ2BP and other RBPs did not mislocalize RPS-2 from mitochondria (not shown), the identities of molecules that mediate ribosome-mitochondria coordination with MIRO-1 remain to be determined. We note that an alternative model, in which MIRO-1 specifically mediates the transport of ribosome-positive mitochondria while ribosome-negative mitochondria are transported by other mechanisms, cannot be ruled out by our data. However, we consider it less likely given MIRO-1’s known function in transport and in recruiting other proteins to the mitochondria surface (Guillén-Samander et al., 2021).

## METHODS

### Nematode husbandry and breeding

Caenorhabditis elegans strains were cultured on standard nematode growth medium plates seeded with OP50 *E. coli* bacteria as previously described (Brenner, 1974). Strains were maintained at 20°C for at least two generations post-starvation before imaging. The Bristol N2 strain was used as wildtype and for genetic crosses. All mutant strains were genotyped by PCR and gel electrophoresis or by Sanger sequencing. Unless otherwise stated, all animals shown in this study are larval stage L4. All strains used in this study are listed in Table S1.

### Plasmid cloning

Plasmids were either generated using traditional restriction enzyme-based ligation techniques or by Gibson assembly of overlapping DNA fragments. All plasmids generated in this study are freely available upon request. Genomic DNA sequences were obtained from WormBase (Sternberg et al., 2024). Ribosome and organelle reporter constructs were generated by PCR amplification of the marker gene from N2 genomic DNA followed by in-frame assembly with eGFP (F64L, S65T), TagRFP-T, or mTagBFP2 in the pSM vector. All three fluorophores used are codon-optimized for *C. elegans* and contain synthetic introns. To generate the mitochondrial localization sequence (MLS) for mitoRFP, two complementary oligonucleotides encoding a codon-optimized sequence for the 24aa MLS of chicken mitochondrial aspartate aminotransferase (Jaussi et al., 1985) were annealed before Gibson assembly with TagRFP-T. To generate the TOMM-20TMD::Linker::KLP-16(T128-N587) “MitoTruck” construct, the genomic sequence encoding the N-terminal 55aa of TOMM-20 was ordered as a gBlock from IDT, as was a codon-optimized sequence encoding a 42aa non-repetitive flexible linker peptide based on Arslan et al. (2022). Both gBlocks were Gibson assembled with a PCR fragment of the *klp-16* gene amplified from N2 genomic DNA.

### Generation of transgenic nematode strains

Transgenic strains were generated by injection of plasmid DNA directly into the gonads of young adult animals according to standard protocols (Mello and Fire, 1995). Plasmids for *odr-1p::TagRFP-T*, *elt-7p::TagRFP-T::NLS* or *elt-7p::eGFP::NLS* were used as coinjection markers, and injection mixes were standardized to a concentration of 100ng/µL by addition of empty pSM vector. After injection, F1 progeny expressing the coinjection marker were singled, and F2 progeny were visualized on a compound microscope to identify strains with desired expression levels. Integrated arrays were generated using the TMP/UV method and outcrossed at least seven times before use.

### Random mutagenesis and gene mapping

L4 stage worms expressing wyIs418 were treated with ethyl methanesulfonate as previously described (Brenner, 1974). F1 progeny were picked to three worms per plate, and subsequent F2 progeny were screened on a fluorescent compound microscope for defects in dendritic ribosome positioning. After visual isolation, mutant worms were clonally propagated and outcrossed to N2. The causal variants in each mutant were mapped by whole-genome sequencing and SNV linkage mapping with the MiModD software package run in a Galaxy server. Finally, cell-specific rescue (*drp-1*) or complementation testing with another allele (*trak-1*) was used to confirm the identify of each mutant allele.

### Fluorescence microscopy and sample preparation

All fluorescent imaging was conducted at room temperature with 405, 488 and 561 nm laser lines and an HC PL APO 63x/1.40NA OIL CS2 objective on a Laser Safe DMi8 inverted microscope (Leica) equipped with a VT-iSIM system (BioVision). Images were acquired with an ORCA-Flash4.0 camera (Hamamatsu) controlled by MetaMorph Advanced Confocal Acquisition Software Package. For consistency across experiments, the same settings for laser power and exposure time were used throughout this study.

For steady-state imaging, L4 stage animals were mounted on a 2% agarose pad and immobilized by mounting with 10 mM levamisole dissolved in M9 buffer. Z-stacks of the entire cell were collected. Each channel was acquired together before advancing to the next Z-position. Slides were only imaged within the first 70 min after mounting.

For timelapse imaging, L4 stage animals were gently handled with a hair pick and incubated for ~45 min in 0.5 mM levamisole before mounting in M9 without paralytics on a 10% agarose pad. Images at three Z-positions spanning 1 µm of depth were collected once every 10 sec (RPS-2::GFP and mitoRFP) or every 5 sec (GFP::RAB-7 and mitoRFP) for a total of 10 min. Maximum intensity projections were used in post-processing to make one frame per time point. Slides were only imaged within the first 70 min after mounting.

### Electron microscopy and sample preparation

Transmission electron micrographs were acquired with an SIS Morada CCD camera on an G2 Spirit BioTWIN microscope (FEI Tecnai). Samples were prepared as previously described (Kennerdell et al., 2009; Saalfeld et al., 2012). Briefly, worms were subjected to high-pressure freezing in M9 buffer in a Leica EM HPM100 before freeze substitution in a Leica EM AFS2 and embedding in pure epon. Embedded worms were sectioned on a Leica EM UC7. Serial cross sections 55 nm thick were collected beginning from the nose of the worm.

### Fluorescence quantification and image analysis

Raw images were processed and analyzed using Fiji/ImageJ (Schindelin et al., 2012). All quantifications were done on raw images. Unless otherwise stated, images displayed in this manuscript are not altered beyond linear brightness and contrast adjustments.

For steady-state images, Z-stacks were made into maximum intensity projections. Dendritic puncta numbers of RPS-2::GFP, mitochondria, and organelle markers were manually scored using the ‘multi-point’ tool. To quantify the redistribution of signal in *drp-1* mutants, background signal was removed with a 500 px rolling ball and dendrites were straightened and cropped to 20 px wide. ‘Plot Profile’ was used to measure fluorescence intensity across the length of the dendrite. Finally, the sum of the fluorescent signal in the first 10 µm of the dendrite was divided by the sum of the fluorescent signal in the first 80 µm of the dendrite. To generate linescans of the dendrite and soma, a 1 px line was drawn across the image and ‘Plot Profile’ was used to measure fluorescent signal across the image. To quantify colocalization between RPS-2::GFP and mitochondria, dendrites were straightened and cropped to 20 px wide, or a region of interest was manually drawn around the cell body. The Pearson’s Correlation Coefficient was then quantified using the ‘Colocalization Threshold’ plugin from the ‘GDSC’ package. To quantify the redistribution of signal in MitoTruck experiments, the dendrite was traced with a 35 px line, or a region of interest was manually drawn around the cell body. Background signal was removed with a 500 px rolling ball, and total fluorescence in the region of interest was measured. Finally, the fluorescent signal in the dendrite was divided by the fluorescent signal in the cell body. To score co-occurrence between RAB-5 and RPS-2, LMP-2 and RPS-2, RAB-7 and RPS-2, and RAB-7 and mitochondria, puncta were manually scored for the presence or absence of an associated punctum in the other channel.

To score the dissociation of RAB-7 from mitochondria of *unc-16* mutants in an unbiased manner, RAB-7-negative mitochondria were blindly selected to score for the presence or absence of RPS-2. First, ‘Plot Profile’ was used to measure background BFP::RAB-7 signal in an area of the dendrite devoid of mitochondria, and this background level was subsequently used as a threshold. Mitochondria with BFP::RAB-7 signal above this threshold were considered to be enriched with RAB-7, while mitochondria lacking BFP::RAB-7 signal above background levels were considered to lack RAB-7 enrichment. Of the mitochondria lacking an enrichment of BFP::RAB-7 signal, a subset was randomly selected to score for the presence or absence of RPS-2. RPS-2 was considered to be present at the mitochondrion if its fluorescent signal was also above background levels.

To analyze timelapse imaging data, dendrites were straightened and cropped to 20 px wide. The ‘KymoResliceWide’ plugin was then used to generate a kymograph. Kymographs were scaled by a factor of 4 in the y-axis to enhance the visibility of each frame. Mitochondria movements were manually scored for the presence or absence of associated RPS-2, and RPS-2 movements were likewise scored for the presence or absence of mitochondria. RAB-7 movements were quantified in the same way. To quantify mitochondrion length, a line was manually drawn over each mitochondrion, and measurements were sorted based on motility and presence of RPS-2.

### Electron microscopy analysis

For both the wildtype and *miro-1; mtx-2* datasets, the region just past the base of the sensory cilia of the amphid dendrites was identified, and the next 36 sections were selected for analysis. We followed the twelve amphid dendrites of the left amphid bundle through all 36 sections, corresponding to a depth of ~2 µm (36 sections * 55 nm per section). In each 55 nm section, the number of mitochondria profiles and polysomes in the entire left amphid bundle were scored. Mitochondria were clearly distinguishable from other organelles by their double membrane and dark interior.

We observed many small, electron-dense particles in the dendrites that appeared to be monosomes; however, close examination revealed that such structures were present in multiple consecutive serial sections. As ribosomes are only 20 nm in diameter, it is not possible for a ribosome profile to be visible in multiple consecutive sections that are each 55 nm thick. Therefore, such structures likely represent neurofilaments.

Based on this observation, we set a strict threshold for scoring ribosomes in our EM dataset. We chose to ignore any monosome-like structures and only score polysomes, which are clearly identifiable by their circular “rosette” formation (Palade, 1955), and we confirmed the identify of polysomes by their absence from consecutive serial sections.

### Statistical analysis

All statistical analyses were performed in GraphPad Prism (version 10). All sample sizes, tests and statistical results are stated in the figure legends. When appropriate, a non-parametric test was used to compare non-normally distributed data.

## ACKNOWLEDGEMENTS

We would like to thank members of the Yogev and Hammarlund labs for helpful discussion and advice. We thank Benjamin Clark for expert technical assistance with HPF, which was conducted at the Yale Neuroscience Electron Microscopy Core Facility, and Zhongyuan Zuo for assistance with FS, sample embedding, and serial sectioning, which were conducted at the Yale Center for Cellular and Molecular Imaging. We are grateful to Rick Fetter and Ben Mulcahy for excellent advice on EM analysis and to Mei Zhen for generously providing access to her published EM data (Witvliet et al., 2021). We also thank Reto Gassmann for sharing the *unc-16(prt183)* strain and Kang Shen for sharing the *dli-1(wy50235)*, *trak-1(wy50182)*, and *mtx-2(wy50266)* strains. Some strains were provided by the CGC, which is funded by NIH Office of Research Infrastructure Programs (P40 OD010440). Some illustrations were created using Biorender.com. Whole-genome sequencing was conducted at the Yale Center for Genome Analysis, which is supported by the National Institute of General Medical Sciences of the National Institutes of Health under Award Number 1S10OD030363-01A1. We also thank the Yale Center for Research Computing for the use of research computing infrastructure. This work was supported by NS114400 to SY and NSF Grant No. DGE1752134 to CJR.

---

**Figure S1.**
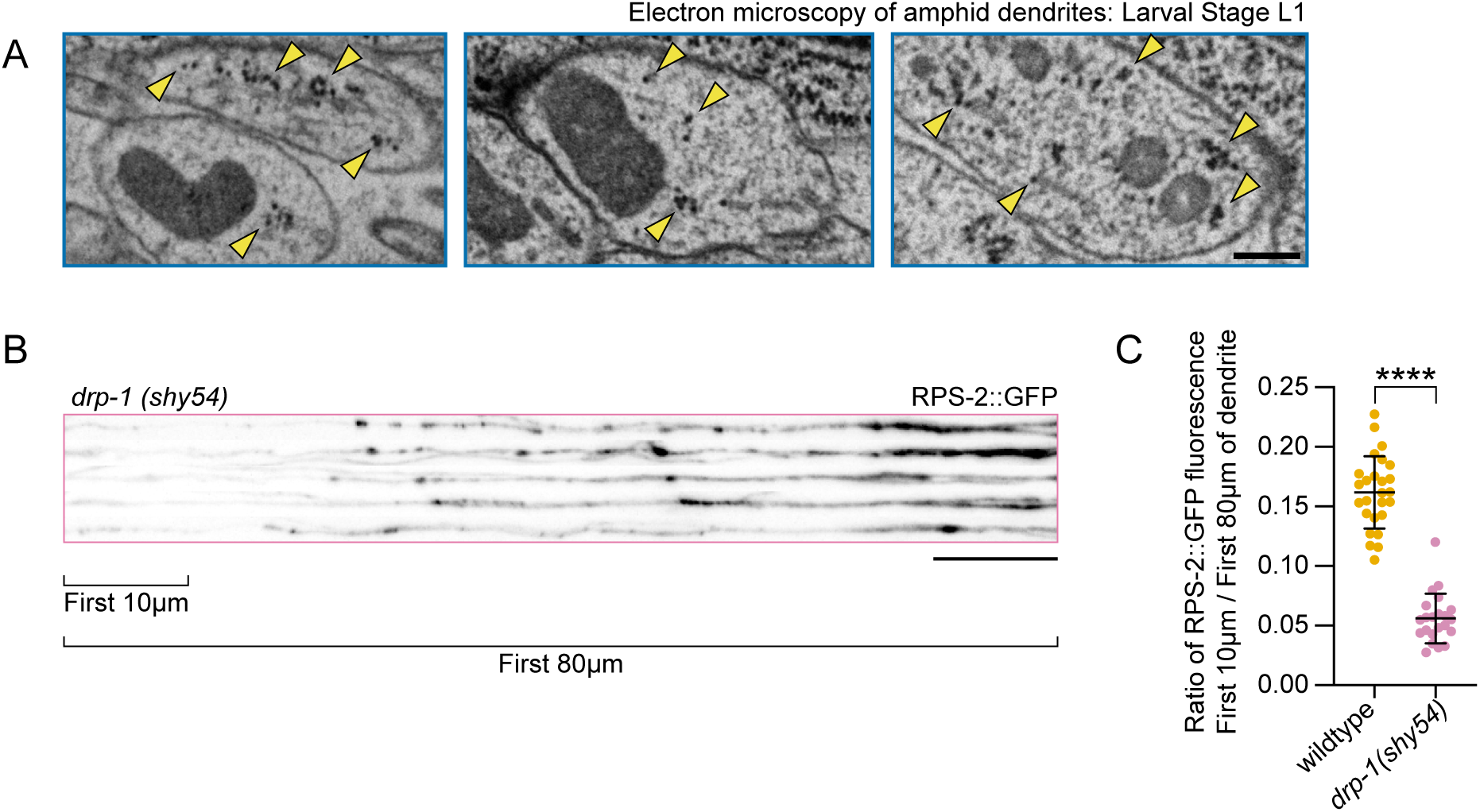
**(A)** Representative electron micrographs of cross sections through the amphid bundle. Yellow arrowheads indicate ribosomes. Data from Witvliet et al. (2021). Scale bar, 200 nm. **(B)** Representative images of RPS-2::GFP expression in *drp-1(shy54)* mutants. A stack of five representative dendrites is shown. Dendrites are straightened and cropped to the first 80 µm of the dendrite beginning from the base of the cilium. Scale bar, 10 µm. **(C)** Quantification of (B). Ratio of RPS-2::GFP fluorescent signal in the first 10 µm of the dendrite normalized to the signal in the first 80 µm of the dendrite, wildtype versus *drp-1(shy54)* mutants. Student’s t-test; n = 26 and 22 animals. Bars indicate mean and SD. The wildtype animals quantified here are the same as those quantified in (Fig. 1 E). ∗∗∗∗p < 0.0001

**Figure S2.**
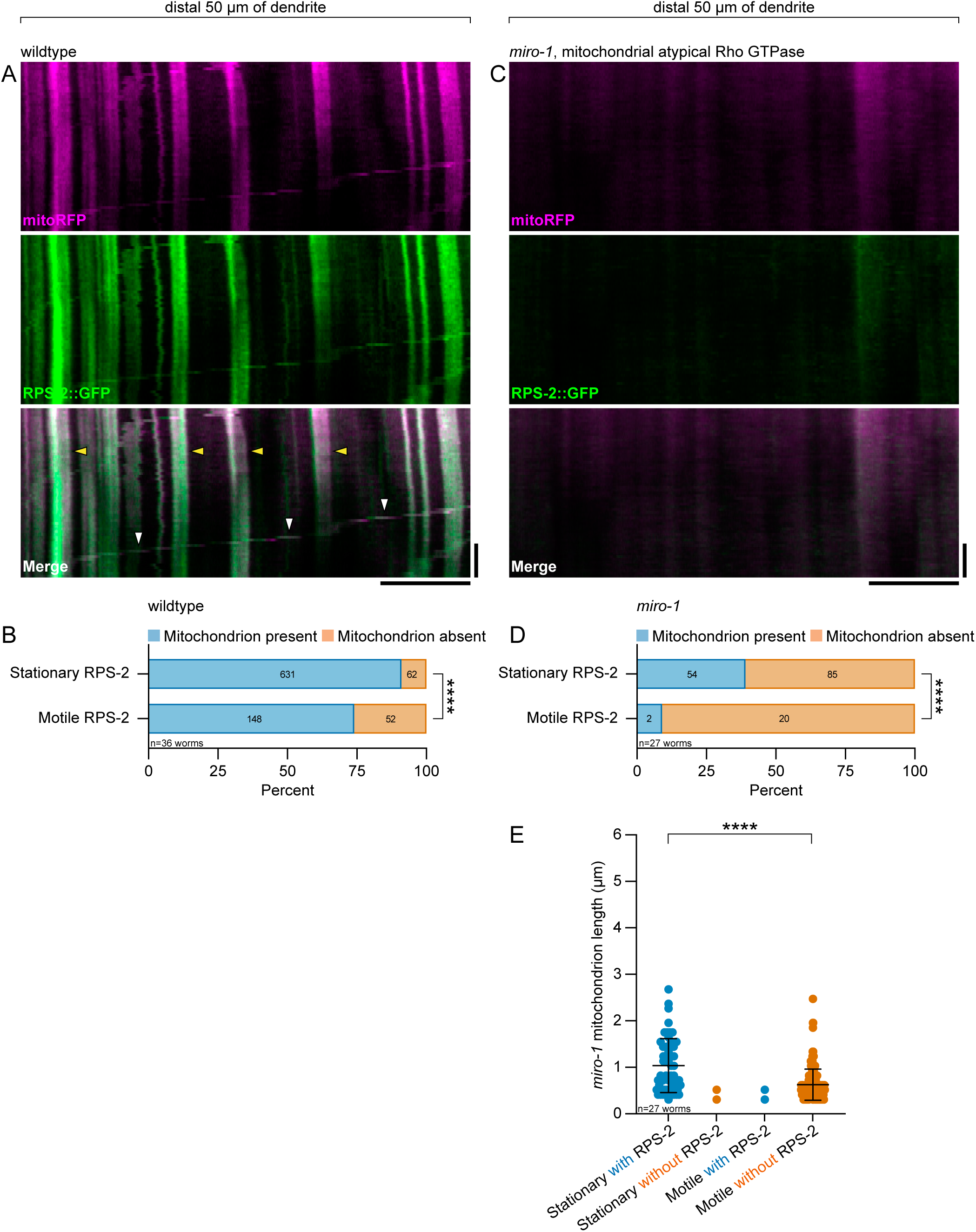
**(A)** Representative kymographs of mitoRFP and RPS-2::GFP movements in the distal dendrite, generated from a 10 min movie with a 10 sec framerate. White arrowheads indicate examples of co-transport between RPS-2::GFP and mitoRFP. Yellow arrowheads indicate examples of stationary mitochondria. Horizontal scale bar, 10 µm; vertical scale bar, 2 min. **(B)** Fisher’s exact test comparing the relationship between motility and mitochondrion association amongst RPS-2 puncta (two-tailed p <0.0001). **(C)** Representative kymographs of dendritic mitoRFP and RPS-2::GFP movement in the distal dendrite of a *miro-1* mutant background. Kymographs are generated and cropped as in (A). Horizontal scale bar, 10µm; vertical scale bar, 2 min. Quantification of wildtype mitochondrion length grouped by motility and presence of RPS-2. n = 631, null, 146, and 123 mitochondria from 36 animals. One-way ANOVA with Tukey’s post-hoc test. Bars indicate mean and SD. **(D)** Fisher’s exact test as in (B). **(E)** Quantification of mitochondrion length in *miro-1* mutants grouped by motility and presence of RPS-2.

**Figure S3.**
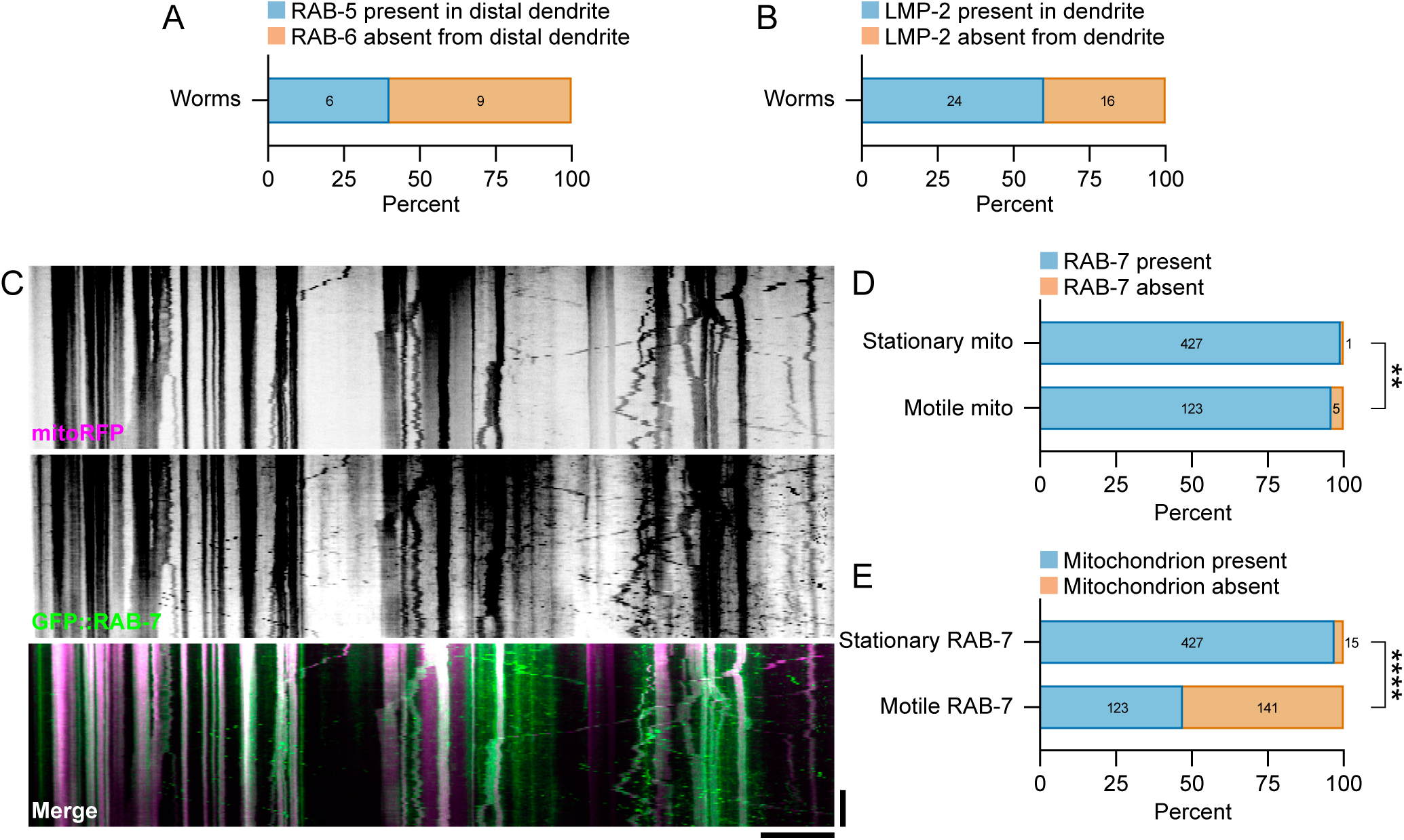
**(A)** Percentage of animals with and without RAB-5 localization in the distal dendrite. **(B)** Percentage of animals with and without LMP-2 localization in the dendrite. **(C)** Representative kymographs of mitoRFP and GFP::RAB-7 in the dendrite, generated from a 10 min movie with a 5 sec framerate. **(D and E)** Quantification of C. The percentage of stationary or motile mitochondria with or without associated RAB-7 (D), and the percentage of stationary or motile RAB-7 signals with or without associated mitochondria. Fisher’s exact tests were used to compare the relationship between signal association and motility. ∗∗∗∗p < 0.0001; ∗∗p < 0.01

**Figure S4.**
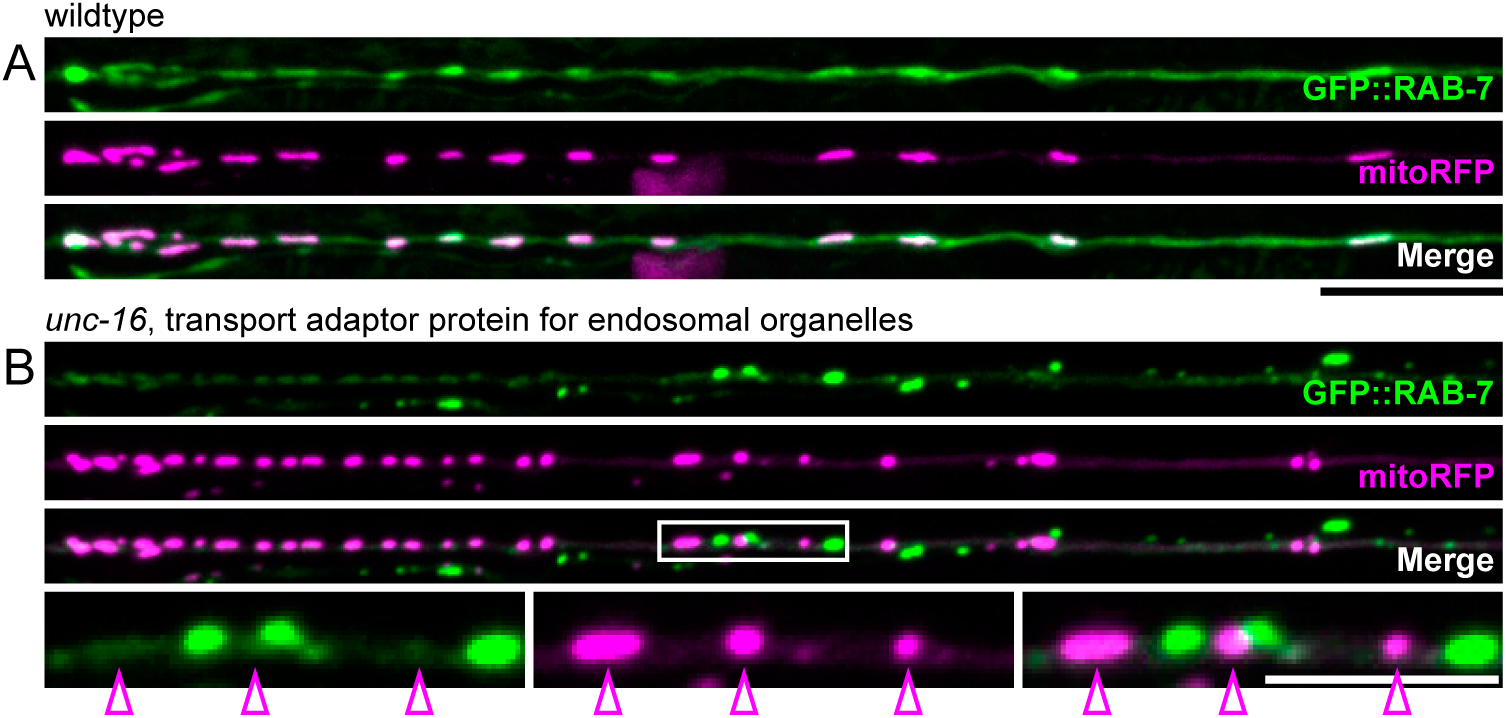
**(A)** GFP::RAB-7 and mitoRFP localization in a wildtype dendrite. **(B)** GFP::RAB-7 and mitoRFP localization in an *unc-16* mutant dendrite. White box indicates region of inset. Inset: arrowheads highlight examples of dissociation between RAB-7 and mitochondria in *unc-16* mutants.

